# RT-qPCR analyses on the osteogenic differentiation from human iPS cells: An investigation of reference genes

**DOI:** 10.1101/2020.05.27.118000

**Authors:** Kensuke Okamura, Yusuke Inagaki, Takeshi K. Matsui, Masaya Matsubayashi, Tomoya Komeda, Munehiro Ogawa, Eiichiro Mori, Yasuhito Tanaka

## Abstract

Reverse transcription quantitative PCR (RT-qPCR) is used to quantify gene expression and require standardization with reference genes. We sought to identify the reference genes best suited for experiments that induce osteogenic differentiation from human induced pluripotent stem (iPS) cells. They were cultured in an undifferentiated maintenance medium and after confluence, further cultured in an osteogenic differentiation medium for 28 days. RT-qPCR was performed on undifferentiation markers, osteoblast and osteocyte differentiation markers, and reference gene candidates. The expression stability of each reference gene candidate was ranked using four algorithms. General rankings identified TATA box binding protein (TBP) in the first place, followed by transferrin receptor (TFRC), ribosomal protein large P0 (RPLP0), and finally, beta-2-microglobulin (B2M), which was revealed as the least stable. Interestingly, universally used GAPDH and ACTB were found to be unsuitable. Our findings strongly suggest a need to evaluate the expression stability of reference gene candidates for each experiment.

---

Quantitative real-time PCR (qPCR) was developed in the 1990s^1^ and is now widely used in various fields as a tool for the accurate and easy detection as well as quantification of target nucleic acid samples. In particular, reverse transcription quantitative PCR (RT-qPCR) is regarded as an indispensable experimental technique for analysis of gene expression by quantifying RNA.

RT-qPCR is used to measure the amplification of an exponentially amplified region and quantify its initial template cDNA, that is, mRNA. Accurate assessments require the correction of inter sample variations, particularly of factors such as the total amount and quality of RNA, reverse transcription efficiency, and PCR efficiency. To perform this, the internal control genes are selected as reference genes, which are quantified along with the target gene. Standardization is performed by calculating their expression ratios. The “housekeeping genes”, those with constant expression levels in the tissues or cells to be analyzed, are generally used as internal control genes; however, their expression levels may vary each tissue and each cell. Fluctuations have been reported depending on the conditions such as the stages of development^2^. The Minimum Information for Publication of Quantitative Real-Time PCR Experiments (MIQE), a set of guidelines compiled to ensure the reproducibility of qPCR, stipulate that reference genes suited for each cell, tissue, and experimental design be selected from numerous candidates so as to ensure reliability of normalisation^3^. Therefore it is evident that, the selection of suitable reference genes, along with the provision of data demonstrating their expression stability, is becoming an indispensable part of quantitative analysis of gene expression using RT-qPCR.

To date, research has mainly focused on the osteogenic potential of mesenchymal stem cells (MSCs)^4,5^. However, since Takahashi and Yamanaka^6^ reported about human induced pluripotent stem (iPS) cells, there have been high expectations for the use of human iPS cells in areas such as regenerative medicine^7^ and the development of drugs for rare diseases^8^. Cell culture experiments have focused on hypoxic environments as the original physiological state inside living tissues^9–11^. We reported that when rat MSCs cultured in a hypoxic environment were exposed to a normoxic environment, it enhanced their osteogenic potential^12^. Based on these results, we expect an examination of the osteogenic differentiation cultures of iPS cells under hypoxic conditions would significantly contribute to osteochondral regenerative medicine and our understanding of the pathology of musculoskeletal diseases such as knee osteoarthritis. Nevertheless, only a few studies have thoroughly verified the expression stability of reference genes used during RT-qPCR for osteogenic differentiation experiments using stem cells, especially iPS cells. Therefore, in the present study we compared multiple internal control genes to identify the most stable reference genes for osteogenic differentiation experiments using RT-qPCR on human iPS cells under various oxygen concentrations.

## Results

### Tissue staining

Continuous induction of osteogenic differentiation (Figure 1a) resulted in areas that revealed calcification (Figure 1b). Samples cultured on day 28 were stained to verify this. The results presented areas positive for Alizarin Red S throughout the wells, with certain regions exhibiting ALP activity (Figure 1c).

**Figure 1.**
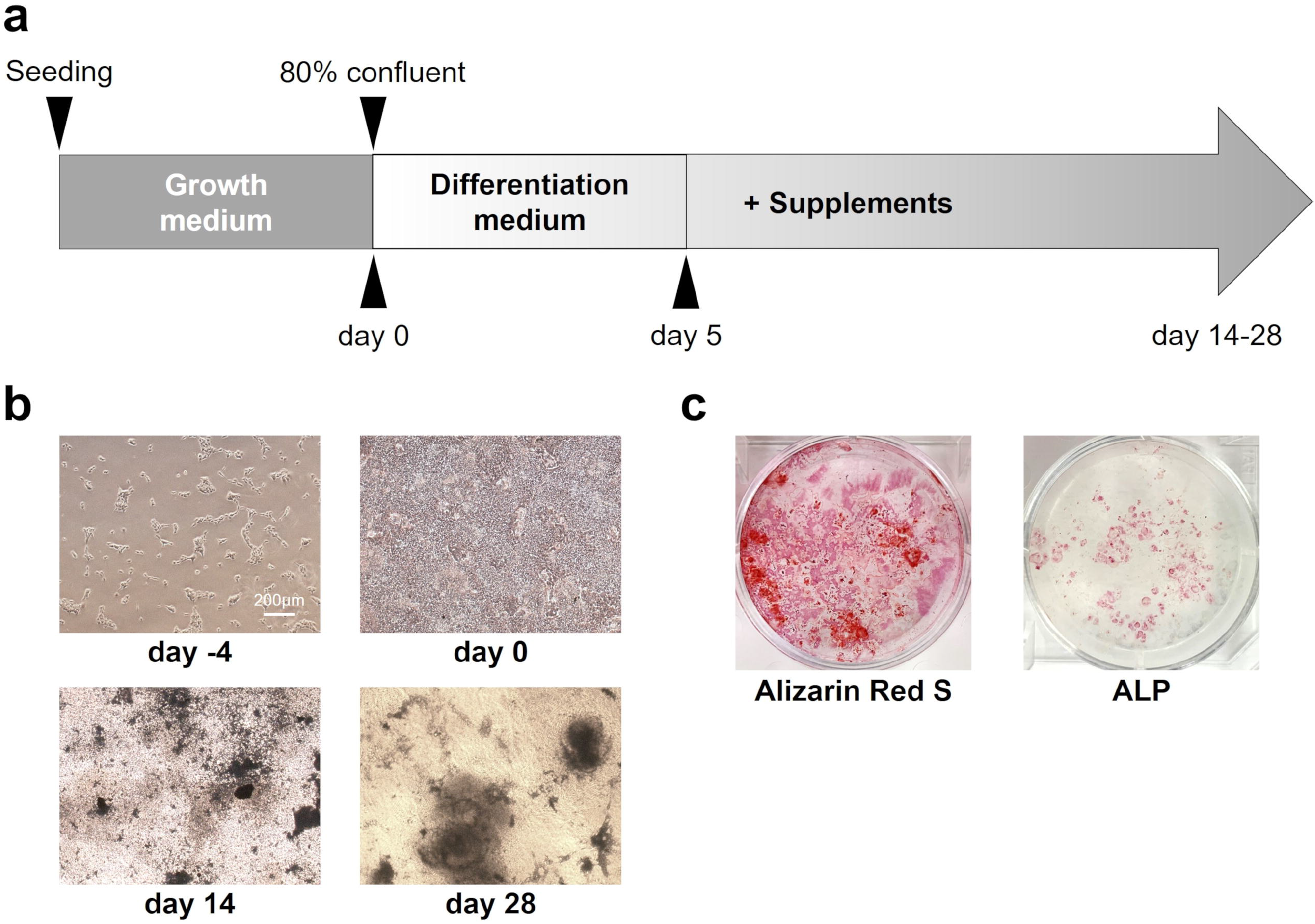
Osteogenic differentiation in human induced pluripotent stem (iPS) cells. (a) Schema of the osteogenic differentiation protocol. (b) Microscopic images on day −4, 0, 7, 14 and 28 showed mineralization yield. (c) Mineralization on day 28 with Alizarin red S and alkaline phosphatase (ALP) staining results in six-well culture plates.

### Comparison of reference gene candidate stability by RT-qPCR

RT-qPCR was performed to compare the expression stability of internal control genes listed in Table 1 on samples from three wells on days 0, 7, 14, 21, and 28. The resulting threshold cycle (Ct) values (Figure 2) and PCR efficiency values from the validation data of each primer were used in each method to calculate the results listed in Table 2. None of the samples were excluded as invalid while verifying RNA purity with absorbance measurements, and all Ct values were < 35.

**Table 1.** Primers of candidate reference genes.

| Symbol | Name | Function | Assay ID<br>(Bio-Rad) | Amplicon<br>length (bp) | Product<br>efficiency |
| --- | --- | --- | --- | --- | --- |
| ACTB | actin, beta | Cytoskeletal structural protein | qHsaCED0036269 | 62 | 2.03 |
| B2M | beta-2-microglobulin | Beta chain of MHC class I | qHsaCID0015347 | 123 | 1.98 |
| G6PD | glucose-6-phosphate dehydrogenase | Maintain the level of NADPH | qHsaCED0001353 | 137 | 2.05 |
| GAPDH | glyceraldehyde-3-phosphate dehydrogenase | Glycolysis and gluconeogenesis | qHsaCED0038674 | 117 | 1.97 |
| GUSB | glucuronidase, beta | Carbohydrate metabolic process | qHsaCID0011706 | 79 | 2.03 |
| HMBS | hydroxymethylbilane synthase | Heme biosynthetic process | qHsaCID0038839 | 137 | 1.98 |
| HPRT1 | hypoxanthine phosphoribosyltransferase 1 | Purine salvage pathway | qHsaCID0016375 | 90 | 2.05 |
| PGK1 | phosphoglycerate kinase 1 | Glycolysis and gluconeogenesis | qHsaCED0003721 | 74 | 1.81 |
| RPL13A | ribosomal protein L13a | Component of the 60S ribosomal subunit | qHsaCED0020417 | 118 | 1.99 |
| RPLP0 | ribosomal protein, large, P0 | Component of the 60S ribosomal subunit | qHsaCED0038653 | 63 | 1.95 |
| RPS18 | ribosomal protein S18 | Component of the 40S ribosomal subunit | qHsaCED0037454 | 67 | 1.94 |
| TBP | TATA box binding protein | Transcription factor binding | qHsaCID0007122 | 120 | 2.01 |
| TFRC | transferrin receptor | Transferrin transport | qHsaCID0022106 | 102 | 1.96 |
| YWHAZ | 14-3-3 protein, zeta polypeptide | Major regulator of apoptotic pathways | qHsaCID0013897 | 175 | 1.96 |

**Figure 2.**
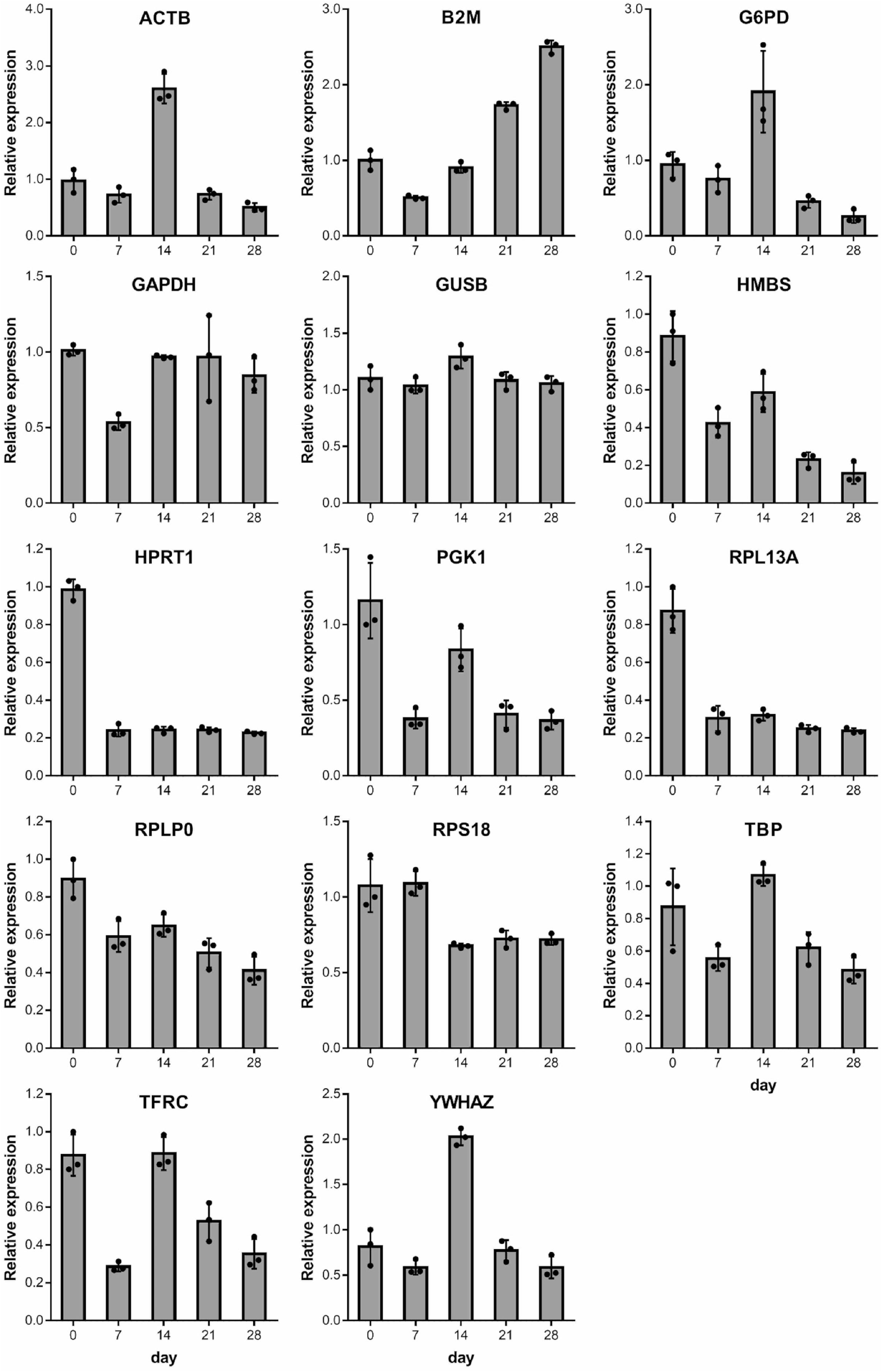
Expression of reference genes in iPSCs during osteogenic differentiation. The values are represented as the mean ± S.D. from three independent samples.

**Table 2.**
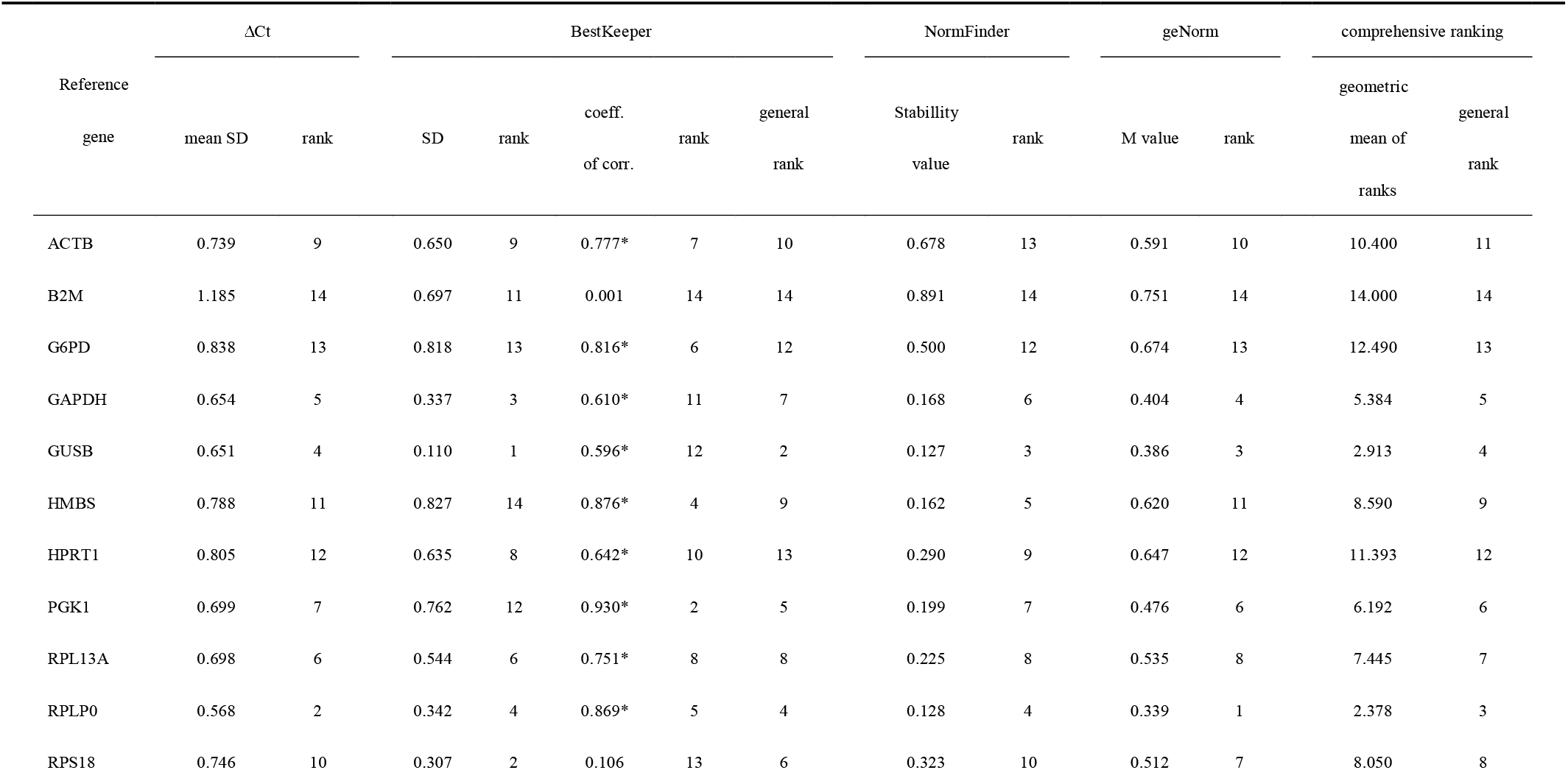
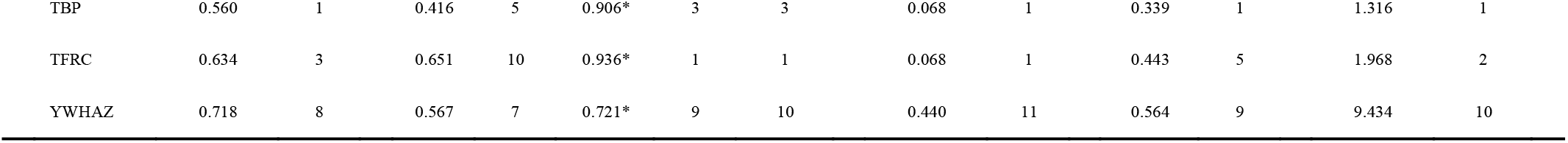
Expression stability and ranking of reference genes, evaluated based on ΔCt method, BestKeeper, NormFinder, and geNorm. *p < 0.05.

First, the ΔCT method revealed that the gene with the lowest mean SD was TBP, and that with the highest value was B2M, which was the only one with a mean SD >1. Based on BestKeeper threshold of SD = 1, no candidate genes were excluded. GUSB and HMBS had the lowest and highest SDs, respectively. In addition, r value was the highest and lowest for TFRC and B2M, respectively. The general ranking calculated from these two rankings revealed that TFRC had the highest stability, while that of B2M was the lowest. Using NormFinder, all the iPS cell samples in this study that underwent osteogenic differentiation were treated as a single group. Three samples from each day were used to analyze a total of 15 samples. The genes with the lowest stability values were TBP and TFRC, whereas B2M had the highest value. Figure 3 depicts two graphs created using geNorm. The pair with the lowest M value was RPLP0 and TBP with threshold values of less than 0.5, whereas B2M presented the highest value (Figure 3a). In addition, V_2/3_-V_13/14_ were all below 0.15 (Figure 3b). These results indicate that valid assessments can be performed with the combination of RPLP0 and TBP. While these methods found several candidates with high stability, including TBP and TFRC, all the methods demonstrated the low stability of B2M.

**Figure 3.**
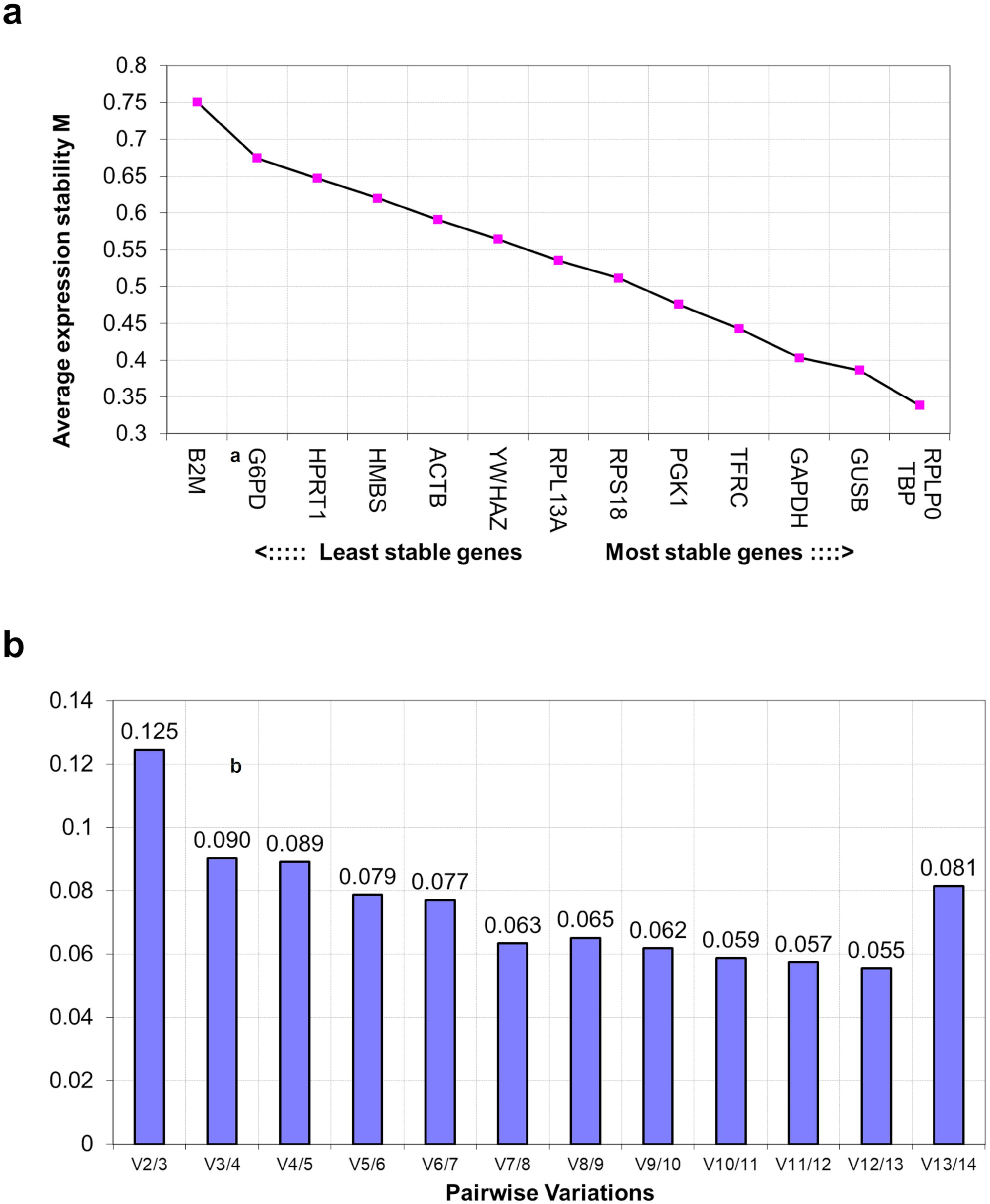
geNorm analysis results. (a) Ranking based on average expression stability M of internal control genes. (b) Optimal number of control genes required for reliable normalization calculated by the pairwise variation (Vn/n+1).

A general ranking was calculated using the geometric means of the rankings in each of the four methods (Table 2). The results reby indicate that TBP is the gene with the most stable expression and that B2M is the lowest.

### Expression of target genes using RT-qPCR

Using TBP as the reference gene based on the results of the aforementioned analysis, RT-qPCR was performed to evaluate the expression of each target gene in the osteogenic differentiation process using the primers listed in Table 3. Similar to the internal control genes, samples from days 0, 7, 14, 21, and 28 were used. To consider the variation in differentiation rates between samples on days 21 and 28, six wells were sampled on these days and three wells were sampled on the other days. For the undifferentiation markers, sampling was performed on days 0, 1, 2, 3, 4, 5, 7, 9, 11, and 14 given that gene expression declines early in the culture period. Due to the variation in differentiation rates between samples, six wells were sampled on days 11 and 14, and three wells were sampled on the other days. Relative expression levels for each gene were calculated using the 2^−^ΔΔCt method by employing the Ct values obtained by RT-qPCR (Figure 4). The verification of RNA purity using absorbance measurements revealed the following samples to be invalid as A260/A230 was greater than 1.8: one sample on day 7 and two samples on day 14 in the undifferentiation marker experiment; one sample on day 14 and one sample on day 21 in the experiment conducted for the other markers. These were excluded from the analysis. As the Ct values of DMP1 exceeded 35 in one sample each on days 7 and 14, these samples were also excluded as they were deemed invalid.

**Table 3.** Primers of target genes

| Assay | Symbol | Name | Function | Assay ID |
| --- | --- | --- | --- | --- |
| PrimePCR<br>SYBR Green Assay<br>(Bio-Rad) | TBP | TATA box binding protein | Transcription factor binding | qHsaCID0007122 |
|  | POU5F1 | POU class 5 homeobox 1 | Marker for undifferentiated cells | qHsaCED0038334 |
|  | NANOG | Nanog homeobox | Marker for undifferentiated cells | qHsaCED0043394 |
|  | ALP | alkaline phosphatase, liver/bone/kidney | Matrix mineralization | qHsaCED0045991 |
|  | OC | osteocalcin | Calcium-binding protein hormone | qHsaCED0038437 |
|  | DMP1 | dentin matrix acidic phosphoprotein 1 | Extracellular matrix mineralization | qHsaCID0015301 |
|  | SOST | sclerostin | Produced primarily by the osteocyte | qHsaCED0046506 |
| Taqman Gene | RUNX2 | runt related transcription factor 2 | Key transcription factor for osteoblast differentiation | Hs00231692_m1 |
| Expression Assay | SP7 | Sp7 transcription factor | Transcription factor for osteoblast differentiation | Hs01866874_s1 |
| (Applied Biosystems) | COL1A1 | collagen type I alpha 1 | Fibrillar collagen | Hs00164004_m1 |

**Figure 4.**
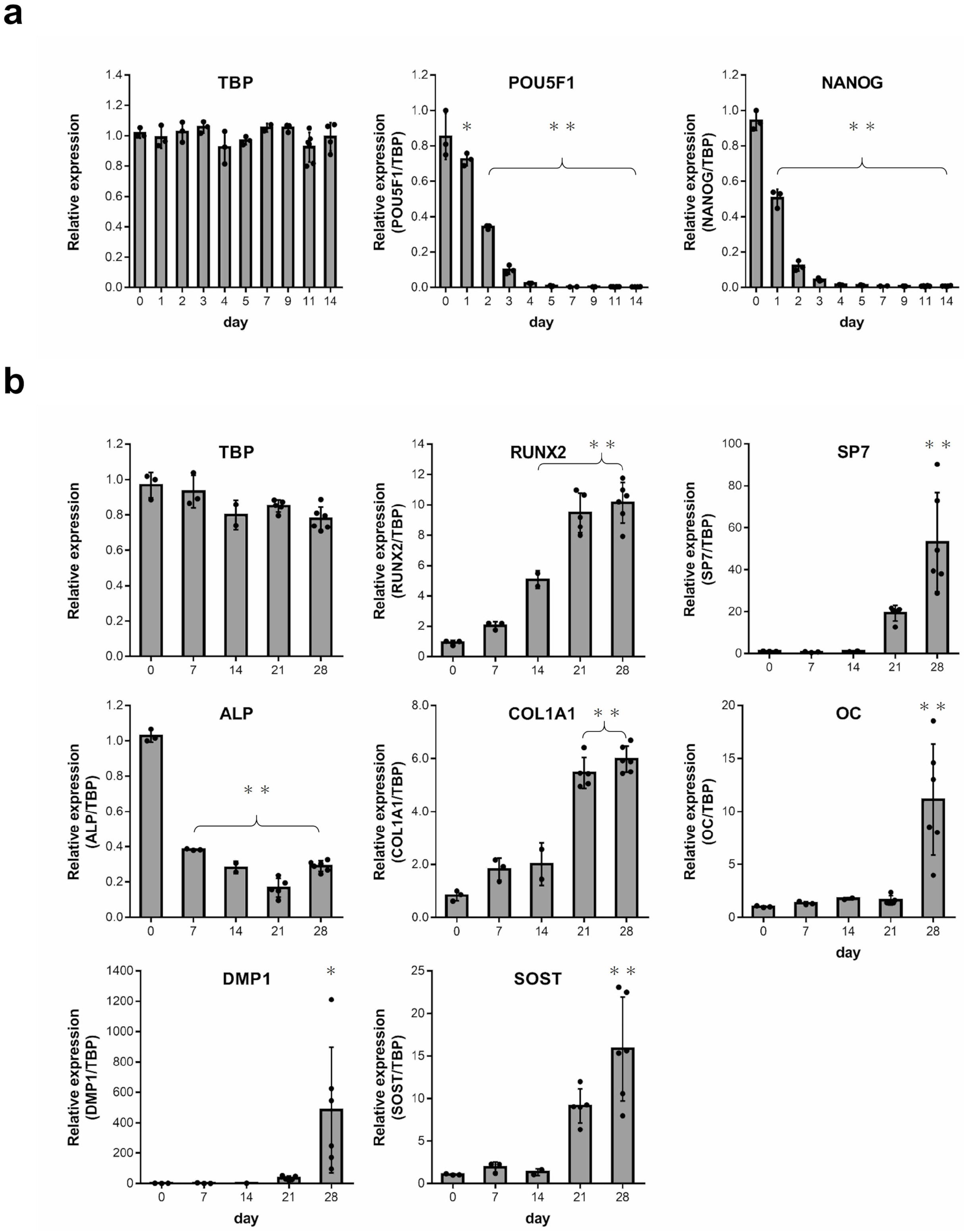
Expression of undifferentiation (a) and osteogenic marker (b) genes in iPSCs during osteogenic differentiation. The values are represented as the mean ± S.D. from two-six independent samples. One-way analysis of variance (ANOVA) with Bonferroni correction for multiple comparisons was applied. *p < 0.05, **p < 0.01 versus day 0.

Figure 4a depicts the expression of TBP as a reference gene and the changes in the relative expression of undifferentiation markers during osteogenic differentiation up to day 14. TBP expression did not vary markedly over the entire period. The expression of the undifferentiation markers, POU5F1 and NANOG, both declined significantly from day 1 until day 14. This suggests that during the osteogenic differentiation process in this study, human iPS cells lost their undifferentiated nature and further differentiated into a certain cell type.

Figure 4b illustrates the expression of TBP as a reference gene and the changes in the relative expression of osteoblast and osteocyte differentiation markers during osteogenic differentiation up to day 28. TBP expression did not significantly vary over the entire period. Expression of the transcription factors, RUNX2 and SP7, and the proteins, COL1A1 and OC, which are osteoblast differentiation markers, exhibited increasing trends starting from day 0. These results indicate that inducing osteogenic differentiation promoted osteoblast differentiation. The protein ALP is not only an osteoblast differentiation marker but also an undifferentiation marker. Its expression level was high on day 0 when the cells were undifferentiated. It then declined over the time, before it exhibited an increasing trend again on day 28 due to osteoblast differentiation. The protein DMP1 is an osteocyte differentiation marker that is expressed in young osteocytes and the protein SOST is expressed in mature bone cells. Both exhibited a marked increase in expression on day 28. Despite the large variations in DMP1 expression between samples on day 28, osteocyte differentiation is assumed to have progressed at least during the second half of the osteogenic differentiation period.

These results indicate that human iPS cells used in the present study differentiated into osteoblasts and osteocytes by inducing osteogenic differentiation. They also provide evidence that assessment of the expression of internal control genes, which was the objective of this study, was performed during the induction of osteogenic differentiation in human iPS cells.

## Discussion

Presently, we conduct osteogenic differentiation experiments using human iPS cells, and RT-qPCR is used as the primary method to quantify gene expression. This speaks to the importance of the accuracy of RT-qPCR and the need to appropriately evaluate the quality of the extracted RNA and select reagents and reference genes.

In the present study, we initially examined the RNA extraction from calcified samples. First, the absorbance of the RNA samples was measured to calculate A_260_/A_230_ and A_260_/A_280_ ratios. High RNA purity was observed when A_260_/A_230_ >1.8 and A_260_/A_280_ >2.0. Denatured agarose gel electrophoresis was also performed on the RNA samples to calculate the ribosomal RNA ratio 28S/18S and determine whether it was close to 2:1. Assessing the RNA quality of several extraction reagents revealed that TRIzol had the highest probability of clearing the RNA purity criteria with 28S/18S close to 2:1. This reagent was found to yield maximum RNA from samples of the same size. Based on these results, we determined that TRIzol is the most suitable for extracting RNA in osteogenic differentiation experiments and devised a method of using RNeasy Micro Kit to recover RNA after extraction.

As described above, Reference Genes H96 comprises preset control wells for verifying genomic DNA contamination, PCR efficiency, RNA quality, and reverse transcription efficiency. We used this plate in a preliminary experiment for the present study and found a case that did not meet the standard for reverse transcription efficiency during control. This prompted us to compare multiple cDNA synthesis kits using reagents used in control wells for the reverse transcription efficiency of Reference Genes H96. The iScript Advanced cDNA Synthesis Kit had the highest reverse transcription efficiency and was adopted for cDNA synthesis.

After this verification process, we verified suitable reference genes, which was the aim of the present study. Standardizing the expression of target genes using reference genes is an essential means of correcting variations between samples in areas such as total RNA, reverse transcription efficiency, and PCR efficiency. The MIQE guidelines state that suitable reference genes should be selected for each tissue, cell, and experimental design^3^. However, it was found that an existing study on osteogenic differentiation using human embryonic stem (ES) cells, MSCs, and iPS cells opted for customarily used genes such as GAPDH, ACTB, and 18S rRNA^13^. The use of GAPDH is particularly common, although no basis has been provided for its selection^14^. Only limited studies have attempted to identify optimal reference genes, and as far as we could find, none for human iPS cells.

To verify the optimal reference genes, the present study used the most common and reliable methods: the ΔCt method, BestKeeper, NormFinder, and geNorm. The ΔCt method is a pairwise comparison, and details on the other three algorithms have been cited in the MIQE guidelines^3^. Reference genes verification results based on these algorithms have been reported in several studies for various cells and differentiation processes^15–23^. For example, Robledo et al.^16^ and Augustyniak et al.^17^ performed verifications using the four algorithms which we used in the present study. Jacob et al.^15^ suggested that verifications should be performed with at least three different algorithms. They found that more than 60% of the published studies on RT-qPCR use two algorithms, NormFinder and geNorm, whereas only 2.6% use all four algorithms. This was based on their review of studies published between 2009 and 2012. Although we presume that more recent studies apply more algorithms, Jacob et al.’s finding highlights the fact that only a few studies have conducted thorough reference gene verification.

Due to the weighting of SD and r values, Bestkeeper may be the least user-friendly out of the four algorithms. As no clear ranking method is provided, some studies have chosen not to use this method^18^. In the present study, after excluding items with SD >1 as invalid, we calculated a general ranking by measuring the geometric mean ranking of SD and r value. A web-based tool called RefFinder (http://www.leonxie.com/referencegene.php) automatically executes BestKeeper, NormFinder, and geNorm algorithms when Ct values are entered, and calculates the stability rankings based on each algorithm, as well as a general ranking. Although numerous studies have used the verification results of RefFinder^15,19–21^, we did not adopt it in the present study because of the following reasons: it does not take into account PCR efficiency, which can affect PCR results^21^; its BestKeeper results only include SD but not r and the method used to determine the general ranking is not stipulated.

Of the 14 preset candidate genes in Reference Genes H96, we found B2M to be the most unsuitable reference gene. This finding was consistent across all the algorithms and the results indicate that B2M has high fluctuations compared to the other candidates. Consistent with Jacob et al.^15^ and Vandesompele et al.’s^24^ recommendation to evaluate combinations of three reference genes, our results found TBP, TFRC, and RPLP0 to be suitable reference gene. The details of our findings have been presented in Table 2.

However, these results were obtained only on the osteogenic differentiation of human iPS cells under normoxic conditions. We plan to perform further experiments under hypoxic conditions and need to exclude candidate genes that fluctuate with changes in oxygen concentration. GAPDH is a typical example of such a gene. Among the three recommended genes, TFRC is involved in the hypoxic response, and as it is presumed to fluctuate depending on the oxygen concentration, its exclusion is highly recommended^25^. Hence, if only one reference gene was to be used, it should be TBP, and if a combination was to be used, geNorm results indicate that the standard should be TBP and RPLP0. The present study found TBP to have the highest stability, a finding which is consistent with that of Rauh et al.^22^ They found TBP to be the most stable reference gene in NormFinder and geNorm in three-dimensional osteogenic differentiation culture experiments of human bone marrow derived MSC. Consistent with their results, we also found TFRC to be the second most stable reference gene. However, instead of TBP or TFRC, their investigation of reference genes in a two-dimensional culture experiment identified completely different genes as the most suitable. Li et al.^23^ found that TBP was one of the least stable genes and B2M was one of the most stable in osteogenic differentiation and adipose differentiation culture experiments of human bone marrow-derived and fetal tissue-derived MSCs. which is contradictory to our findings. This reversed stability ranking of genes has also been previously mentioned by Rauh et al^22^. The selection of reference genes suited to each cell, tissue, and experimental design requires an investigation of the expression stability indicated by reference gene candidates in each experiment.

We would like to highlight that the commonly used GAPDH was not highly ranked; its position was 5 out of 14, whereas ACTB was positioned at 11. Our results indicate that commonly used reference genes such as GAPDH and ACTB were unsuitable in the current experiment and that using them may have led to incorrect interpretations of gene expression levels. This is consistent with De Jonge et al.’s^26^ analysis of data from public samples of 13,629 microarrays performed in various tissues and conditions; they found that of the 13,037 genes, none of the commonly used reference genes—ACTB, GAPDH, HPRT1, B2M— were ranked in the top 50 for stability.

Our study standardized the results of RT-qPCR on target genes in the osteogenic differentiation process using only TBP, which was used to evaluate the expression level of each gene. We found that the levels of undifferentiation markers declined markedly within 2 weeks, indicating a loss of undifferentiation, whereas those of the osteoblast and osteocyte differentiation markers increased within 4 weeks. This indicates that iPS cells differentiate into osteoblasts and osteocytes. This observation was also supported by the results of two types of staining. We were able to achieve the main purpose of this study, that is to verify the reference genes in an osteogenic differentiation experiment on human iPS cells.

## Materials and Methods

### Human iPS Cell line and induction of osteogenic differentiation

The commercially available Cellartis Human iPS Cell Line 12 (Takara Bio Inc., Kusatsu, Japan) was used as the human iPS Cell line. This was seeded on wells coated with the Matrigel basement membrane matrix for feeder-free culturing (Corning Inc., Corning, NY, USA) and then cultured in the undifferentiated maintenance medium Essential 8 (E8; Thermo Fisher Scientific Inc., Waltham, MA, USA). The medium was changed daily, and after reaching confluence (about 80%), the cells were detached using Versene (Thermo Fisher Scientific Inc.) (12 minutes in 37°C) and replated. This passage procedure was repeated.

Cells obtained by cloning in this manner were used in the osteogenic differentiation experiment (Figure 1a). Confluent cells (defined as Day 0) were switched to Dulbecco’s modified Eagle’s medium (DMEM; Nacalai Tesque Inc.) containing fetal bovine serum (FBS, 15%; GE Healthcare Life Sciences, Logan, UT, USA), 1% nonessential amino acids (Sigma-Aldrich Inc., St. Louis, MO, USA), 1% antibiotic-antimycotic mixed stock solution (Nacalai Tesque Inc., Kyoto, Japan), and 100 μM β-mercaptoethanol (Nacalai Tesque Inc.). This medium was changed daily. From day 5, 10 mM β-glycerophosphate (Sigma-Aldrich), 50 μg/ml L-ascorbic acid (Wako Pure Chemical Ind., Osaka, Japan), 10 nM dexamethasone (Sigma-Aldrich), and 50 nM 1,25-(OH)2 vitamin D_3_ (Wako Pure Chemical Ind.) were added to the above-mentioned medium to induce osteogenic differentiation and this culture was changed once every two days.

### Tissue staining

For the staining assessments, the human iPS cells that underwent osteogenic differentiation were fixed with 95% ethanol. To evaluate Ca deposition, they were stained with Alizarin Red S (Nacalai Tesque Inc.). To assess ALP activity, they were stained with Naphthol AS-MX phosphate disodium salt (Sigma-Aldrich) and Fast Red Violet LB Salt (Nacalai Tesque Inc.) in 56 mM 2-amino-2-methyl-1,3-propanediol buffer (Wako Pure Chemical Ind.) adjusted to pH 9.9.

### Gene expression analysis

For total RNA extraction, TRIzol (Invitrogen Life Technologies, Carlsbad, CA, USA) was added and left to stand at room temperature for 10 minutes. The resulting solution was transferred to a QIA shredder spin column (Qiagen Inc., Valencia, CA, USA) and centrifuged at 15,000 rpm for 2 minutes at room temperature. The resulting filtrate was transferred to a new 1.5 ml microtube, 200 μl of chloroform was added, mixed well, and allowed to stand at room temperature for 5 minutes. This was centrifuged at 15,000 rpm for 15 minutes at 4 °C, the upper aqueous layer was transferred to a new 1.5 ml microtube, and the same amount of 70% ethanol was added and mixed well. This sample was transferred to RNeasy Micro Kit spin column (Qiagen Inc.) and then total RNA was recovered according to the procedure described in the product’s instruction manual. Next, DNase treatment was performed to prevent genomic DNA contamination. The resulting solution was diluted 20-fold with Tris-EDTA buffer (pH 8.0) (Nippon Gene, Toyama, Japan), and then analyzed with a Ultrospec 3000 spectrophotometer (Biochrom Ltd., Cambridge, England) at 230, 260, and 280 nm to measure the absorbance, based on which the purity of RNA was confirmed and its concentration was calculated. RNA template (1 μg) was prepared and reverse transcription reaction was performed with iScript Advanced cDNA Synthesis Kit (Bio-Rad Laboratories, Hercules, CA, USA) according to the procedure described in the product’s instruction manual. The StepOnePlus Real-Time PCR System (Applied Biosystems, Foster City, CA, USA) was used for RT-qPCR.

Reference Genes H96 (Bio-Rad Laboratories) was used in RT-qPCR to compare the expression levels of several internal control genes that were candidate reference genes. Reference Genes H96 is a predesigned plate with wells containing primers for 14 types of validated human internal control genes (Table 1), which can be used as control assays for evaluating genomic DNA contamination, PCR efficiency, RNA quality, and reverse transcription efficiency. A reaction solution of 20 Δl comprising 10 μl 2× Sso Advanced universal SYBR Green Supermix (Bio-Rad), 1 μl cDNA template (25 ng), and 9 μl sterile distilled water was added per sample in each well. For the reaction conditions, initial activation was performed at 95 °C for 2 minutes, followed by 40 cycles of thermal denaturation at 95 °C for 5 s and then annealing and elongation at 60 °C for 30 s, and eventually the melting curve was analyzed.

The internal control genes selected based on the data analysis described below were used as the target genes in RT-qPCR to analyze the expression of undifferentiation, osteoblast differentiation, and osteocyte differentiation markers (Table 3). PrimePCR SYBR Green Assay (Bio-Rad) or Taqman Gene Expression Assay (Applied Biosystems) was used as the primer. For the former, 5 μl of 2× SsoAdvanced universal SYBR Green Supermix, 0.5 μl of 20× SYBR Green Assay, 1 μl (25 ng) of cDNA template, and 3.5 μl of sterile distilled water were used as the reaction solution, for a total of 10 μl per sample dispensed in each well. The reaction conditions were the same as for Reference Genes H96. For the latter, 10 μl of 2× Taqman Fast Universal PCR Master Mix, 1 μl of 20× Gene Expression Assay, 2 μl (50 ng) of cDNA template, and 7 μl of sterile distilled water were used as the reaction solution, for a total of 20 Δl per sample dispensed in each well. For the reaction conditions, initial activation was performed at 95 °C for 20 s, followed by 40 cycles of thermal denaturation at 95 °C for 1 s, and annealing and elongation at 60 °C for 20 s.

### Data analysis

To select suitable reference genes for osteogenic differentiation experiments in human iPS cells, Ct values, PCR efficiency, and relative expression values based on the internal control genes obtained using RT-qPCR were used in the following methods for analysis.

### ΔCt method

Silver et al.^27^ reported the ΔCt method for comparing the stability of candidate genes based on pairwise comparison. A pair is created with candidate gene A and another candidate gene B. Thereafter, the difference in their Ct values in the same sample is calculated to obtain ΔCt. The standard deviation (SD) of ΔCt in all samples for the pair of gene A and B is then calculated (A vs B). This calculation is also performed for all the pairs containing gene A such as A vs C and A vs D. The arithmetic means of the all SD (mean SD) for gene A represents the variation in the expression of gene A with respect to all the candidate genes, allowing it to be compared with other genes. Accordingly, the gene with the lowest value is the most stable reference gene. A characteristic of this method is that assessments can be performed by merely repeating a relatively simple calculation.

### BestKeeper

The BestKeeper algorithm of Pfaffl et al.^28^ is characterized by directly inputting the crossing point (CP) value, which is equal to Ct, and PCR efficiency for all the samples of each candidate reference gene to automatically calculate two indicators. The first indicator is the SD of the CP values of each gene, with the lowest value being the most stable reference gene. The developers recommend removing any candidate gene with SD >1. Next, the geometric mean of the CP values of all genes in each sample is calculated as the BestKeeper index, and Pearson’s correlation coefficient (r) between this and the CP value of each gene is measured as the second indicator. The highest value represents the most stable reference gene. Since BestKeeper requires an evaluation of both SD and r, we considered the geometric mean of the ranking for both indicators, which was used to create a general ranking for BestKeeper.

### NormFinder

The NormFinder algorithm of Andersen et al.^29^ is characterized by evaluating stability through the amount of change in expression within and between groups. A “stability value” is obtained by inputting the relative expression value, with the lowest value representing the most stable reference gene. Each group needs at least eight samples for appropriate assessments and having 5-10 candidate reference genes is recommended.

### geNorm

The geNorm algorithm of Vandesompele et al.^24^ is based on pairwise comparisons. Inputting the relative expression values calculates two indicators. The “M value” indicates the stability of expression, with the lowest value representing the most stable reference gene. In the developer’s experimental data, genes with stable expression had a mean M value < 0.5^30^. When evaluating the stability of two or more reference gene combinations, a pairwise variation (V_n/n+1_) is calculated to determine the appropriate number of reference genes to be combined, and V_n/n+1_ < 0.15 is considered as an appropriate cutoff value.

### Statistical analysis

The expression of each gene is presented as mean ± standard deviation (SD). When a significant difference was observed in one-way analysis of variance (ANOVA) with day as the factor, multiple comparisons with the Bonferroni method were performed. P < 0.05 was considered as statistically significant. SPSS 22.0J for Windows (SPSS Inc., Chicago, IL, USA) was used as the statistical software.

## Acknowledgements

This work was supported by grants from JSPS KAKENHI [JP18K1663 and 20K09508 to Y.I., JP20H03199 to E.M.], AMED Brain/MINDS Beyond [JP20dm0307032 to E.M.], Takeda Science Foundation to E.M., Uehara Memorial Foundation to E.M., Naito Foundation to E.M., MSD Life Science Foundation to E.M., and the unrestricted funds provided to E.M. from Dr. Taichi Noda (KTX Corp., Aichi, Japan) and Dr. Yasuhiro Horii (Koseikan, Nara, Japan). The authors thank Keren-Happuch E Fan Fen for critically reading of the manuscript.

## Author contributions

K.O. designed and conducted the experiments, analyzed the data and drafted the manuscript. Y.I. and E.M. designed the study and wrote and revised the manuscript. T.K.M., M.M. and T.K. conducted the experiments. T.K.M. and M.M. supervised the experiments and data analyzing. O.M., E.M. and Y.T. supervised this project. All authors reviewed the manuscript.

## Competing interests

The authors declare no competing interests.

## Notes

### Competing Interest Statement

The authors have declared no competing interest.

## References

1) Higuchi, R., Fockler, C., Dollinger, G., & Watson, R. Kinetic PCR analysis: Real-time monitoring of DNA amplification reactions. Biotechnology (N Y) 11, 1026–1030 (1993).

2) Bustin, S. A. Quantification of mRNA using real-time reverse transcription PCR (RT-PCR): Trends and problems. J Mol Endocrinol 29, 23–39 (2002).

3) Bustin, S. A. et al. The MIQE guidelines: Minimum information for publication of quantitative real-time PCR experiments. Clin Chem 55, 611–622 (2009).

4) Akahane, M. et al. Osteogenic matrix sheet-cell transplantation using osteoblastic cell sheet resulted in bone formation without scaffold at an ectopic site. J Tissue Eng Regen Med 2, 196–201 (2008).

5) Tohma, Y. et al. Bone marrow-derived mesenchymal cells can rescue osteogenic capacity of devitalized autologous bone. J Tissue Eng Regen Med 2, 61–68 (2008).

6) Takahashi, K., & Yamanaka, S. Induction of pluripotent stem cells from mouse embryonic and adult fibroblast cultures by defined factors. Cell 126, 663–676 (2006).

7) Tsumaki, N., Okada, M., & Yamashita, A. iPS cell technologies and cartilage regeneration. Bone 70, 48–54 (2015).

8) Hino, K. et al. Activin-A enhances mTOR signaling to promote aberrant chondrogenesis in fibrodysplasia ossificans progressiva. J Clin Invest 127, 3339–3352 (2017).

9) Grayson, W. L., Zhao, F., Bunnell, B., & Ma, T. Hypoxia enhances proliferation and tissue formation of human mesenchymal stem cells. Biochem Biophys Res Commun 358, 948–953 (2007).

10) Hu, X. et al. Transplantation of hypoxia-preconditioned mesenchymal stem cells improves infarcted heart function via enhanced survival of implanted cells and angiogenesis. J Thorac Cardiovasc Surg 135, 799–808 (2008).

11) Ren, H. et al. Proliferation and differentiation of bone marrow stromal cells under hypoxic conditions. Biochem Biophys Res Commun 347, 12–21 (2006).

12) Inagaki, Y. et al. Modifying oxygen tension affects bone marrow stromal cell osteogenesis for regenerative medicine. World J Stem Cells 9, 98–106 (2017).

13) Yang, X. et al. Bone to pick: the importance of evaluating reference genes for RT-qPCR quantification of gene expression in craniosynostosis and bone-related tissues and cells. BMC Res Notes 5, 222 (2012).

14) Bustin, S. A., & Nolan, T. Data analysis and interpretation in A-Z of Quantitative PCR (ed. Bustin, S. A.) 441–492 (International University Line, 2004).

15) Jacob, F. et al. Careful selection of reference genes is required for reliable performance of RT-qPCR in human normal and cancer cell lines. PLoS One 8, e59180 (2013).

16) Robledo, D. et al. Analysis of qPCR reference gene stability determination methods and a practical approach for efficiency calculation on a turbot (Scophthalmus maximus) gonad dataset. BMC Genomics 15, 648 (2014).

17) Augustyniak, J., Lenart, J., Lipka, G., Stepien, P. P., & Buzanska, L. Reference gene validation via RT-qPCR for human iPSC-derived neural stem cells and neural progenitors. Mol Neurobiol 56, 6820–6832 (2019).

18) Piehler, A. P., Grimholt, R. M., Ovstebø, R., & Berg, J. P. Gene expression results in lipopolysaccharide-stimulated monocytes depend significantly on the choice of reference genes. BMC Immunol 11, 21 (2010).

19) Diesel, L. F. et al. Stability of reference genes during tri-lineage differentiation of human adipose-derived stromal cells. J Stem Cells 10, 225–242 (2015).

20) Taïhi, I. et al. Validation of housekeeping genes to study human gingival stem cells and their in vitro osteogenic differentiation using real-time RT-qPCR. Stem Cells Int 2016, 6261490 (2016).

21) De Spiegelaere, W. et al. Reference gene validation for RT-qPCR, a note on different available software packages. PLoS One 10, e0122515 (2015).

22) Rauh, J., Jacobi, A., & Stiehler, M. Identification of stable reference genes for gene expression analysis of three-dimensional cultivated human bone marrow-derived mesenchymal stromal cells for bone tissue engineering. Tissue Eng Part C Methods 21, 192–206 (2015).

23) Li, X. et al. Identification of optimal reference genes for quantitative PCR studies on human mesenchymal stem cells. Mol Med Rep 11, 1304–1311 (2015).

24) Vandesompele, J. et al. Accurate normalization of real-time quantitative RT-PCR data by geometric averaging of multiple internal control genes. Genome Biol 3, research0034 (2002).

25) Bianchi, L., Tacchini, L., & Cairo, G. HIF-1-mediated activation of transferrin receptor gene transcription by iron chelation. Nucleic Acids Res 27, 4223–4227 (1999).

26) De Jonge, H. J. et al. Evidence based selection of housekeeping genes. PLoS One 2, e898 (2007).

27) Silver, N., Best, S., Jiang, J., & Thein, S. L. Selection of housekeeping genes for gene expression studies in human reticulocytes using real-time PCR. BMC Mol Biol 7, 33 (2006).

28) Pfaffl, M. W., Tichopad, A., Prgomet, C., & Neuvians, T. P. Determination of stable housekeeping genes, differentially regulated target genes and sample integrity: BestKeeper - Excel-based tool using pair-wise correlations. Biotechnol Lett 26, 509–515 (2004).

29) Andersen, C. L., Jensen, J. L., & Ørntoft, T. F. Normalization of real-time quantitative reverse transcription-PCR data: a model-based variance estimation approach to identify genes suited for normalization, applied to bladder and colon cancer data sets. Cancer Res 64, 5245–5250 (2004).

30) Hellemans, J., Mortier, G., De, P. A., Speleman, F., & Vandesompele, J. qBase relative quantification framework and software for management and automated analysis of real-time quantitative PCR data. Genome Biol 8, R19 (2007).

